# Deletion of murine IQGAP1 results in increased mTOR activation and blunted ketogenic response

**DOI:** 10.1101/182469

**Authors:** Hanna Erickson, Sayeepriyadarshini Anakk

## Abstract

IQ motif-containing GTPase Activating Protein 1 (IQGAP1) is a ubiquitously expressed scaffolding protein that integrates signaling from multiple cellular processes including motility, adhesion, and proliferation. Here, we show that IQGAP1 is induced in the liver upon fasting and can also regulate β-oxidation and ketone body synthesis. Utilizing ketogenic diet and pharmacologic activation we identified that hepatic PPARα activity is compromised in *Iqgap1*^*-/-*^mice. Our data show that IQGAP1 interacts with the mechanistic target of rapamycin (mTOR) and IQGAP1 deletion results in enhanced mTOR complex 1 (mTORC1) activation. Conversely, ectopic expression of IQGAP1 in *Iqgap1*^*-/-*^ mice was sufficient to suppress mTORC1 signaling. We also confirmed that modulation of mTORC1 signaling by IQGAP1 is cell autonomous. This increased mTORC1 activation impedes PPARα signaling since mTORC1 inhibition restored a subset of metabolic genes in *Iqgap1*^*-/-*^ mice. Overall, we demonstrate a previously unidentified role for IQGAP1 as an important regulator of mTORC1 activity and long-term ketosis.

## Introduction

The IQ motif-containing GTPase Activating Protein (IQGAP) family of multi-domain scaffolding proteins includes three members, of which IQGAP1 is the only ubiquitously expressed isoform (1-5). IQGAP1 interacts with several binding partners and integrates signals that control diverse cellular processes such as cell-cell adhesion via E-Cadherin (6), cell motility via Cdc42 and Rac1 (7), cell proliferation via the Ras/MAP kinase pathway (8), and membrane receptor signaling via EGF receptor (9). More recently, its role in scaffolding PI-3K/ AKT signaling and subsequent glucose homeostasis has been identified (10, 11). Because IQGAP1 regulates several energy-requiring cellular processes, we hypothesized that IQGAP1 may also coordinate the response to changes in metabolic state.

The liver is a central organ responsible for nutrient metabolism. In the fed state, the liver responds to high insulin levels by promoting *de novo* lipogenesis to store excess calories in adipose depots (12). On the other hand, in the fasted state, the liver breaks down fatty acids to synthesize ketone bodies, which are an effective alternate fuel source. Ketogenesis is crucial to maintain energy levels during fasting. It can also be induced nutritionally with a low-carbohydrate, low protein, high-fat diet known as the ketogenic diet (KD). This process is largely regulated transcriptionally by the nuclear receptor PPARɑ (13-18).

It is has been shown that the major fed state sensor mechanistic target of rapamycin (mTOR) complex 1 (mTORC1) (19, 20) can cross-talk with PPARɑ signaling and restrict its activity to the fasting state (21). mTORC1 includes three-core components: mTOR, Regulatory-associated Protein of mTOR (RAPTOR) and mammalian lethal with Sec13 protein 8 (LS8). This complex regulates protein translation through p70-S6 Kinase 1 (S6K1) and eukaryotic translation initiation factor 4E binding protein 1 (4E-BP1) (22, 23), lipid synthesis through sterol responsive element binding protein (SREBP) (24), and glycolysis through phospho-fructo kinase (25). In addition, mTORC1 suppresses fasting-associated ketogenesis (21) and autophagy (26-28). Inversely, during fasting state, mTORC1 activity is suppressed by AMPK (29). Thus, mTORC1 activity cycles up and down between fed and fasted states to couple anabolic and catabolic processes with nutrient availability.

Here we show that compared to the fed state, hepatic IQGAP1 levels are induced upon fasting, and its deletion impairs the ketogenic response. Since PPARa regulates ketogenesis, we examined its pharmacological activation in the presence and absence of IQGAP1. Compared to WT, IQGAP1-null (*Iqgap1*^*-/-*^) mice show blunted response to PPARɑ activation. Mechanistically, *Iqgap1*^*-/-*^ mice exhibit increased hepatic mTORC1 activity, which in turn inhibits the fasting response. Conversely, reintroducing IQGAP1 to the liver of *Iqgap1*^*-/-*^ mice decreases mTORC1 activity and promotes expression of genes involved in ketogenesis. These data together uncover a novel role for IQGAP1 in controlling ketone body synthesis. Importantly, primary liver cells and the well-established human hepatoma HepG2 cell line recapitulate these results, indicating a cell autonomous role for IQGAP1 in regulating mTORC1 activity. Additionally, inhibition of mTOR activity in *Iqgap1*^*-/-*^ mice with rapamycin reverses some of the misregulated PPARɑ responses. Taken together, these results show that IQGAP1 is necessary to elicit appropriate ketogenic response in the liver.

## Results

### Hepatic IQGAP1 deletion alters the fasting response

IQGAP1 integrates multiple signaling pathways, and its expression level largely determines its function as a scaffold (30). Therefore, we examined whether IQGAP1 coordinates the metabolic response to changes in nutrition status by fasting wild-type (WT) mice for 24 hours and analyzing IQGAP1 expression. We found a significant increase in *Iqgap1* transcript levels in the liver (figure 1A) upon fasting that corroborated well with the robust increase in IQGAP1 protein expression (figure 1B). The increase in IQGAP1 expression was specific to the liver and was not observed in white adipose tissue (WAT) (figure S1A).

**Figure 1.**
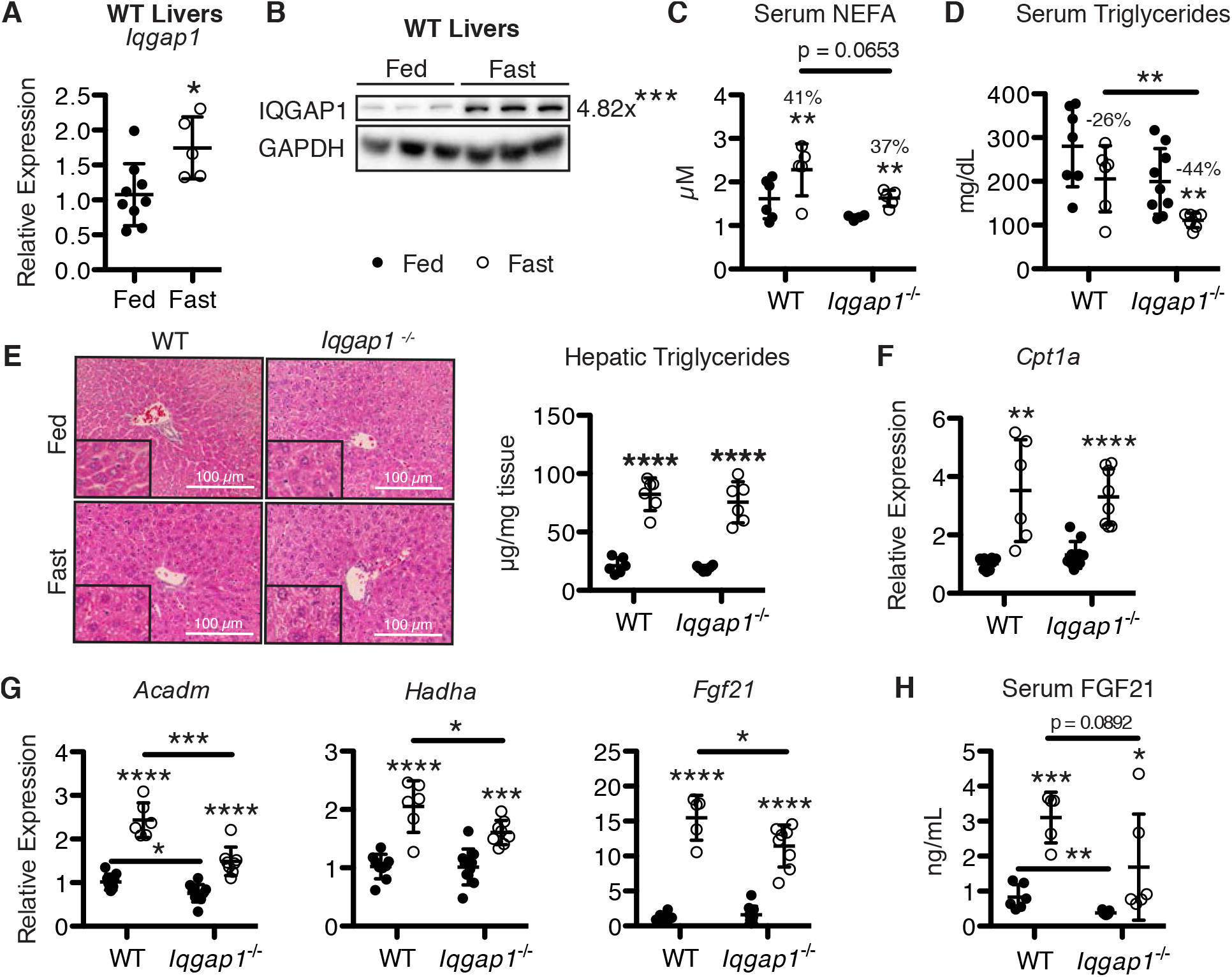
Hepatic IQGAP1 is regulated by fasting and is required for the fasting response. Mice were fed *ad libitum* or fasted for 24 h and sacrificed at ZT 4. (A) *Iqgap1* gene expression normalized to *Gapdh* expression in livers of WT mice as measured by quantitative reverse transcription PCR (qPCR). (B) Immunoblots of liver extracts from WT mice. Each lane consists of a mixture of liver extracts from two animals (n = 6 mice per group). Protein levels were measured by densitometry. IQGAP1 levels were normalized to GAPDH expression. Relative average expression in fasted livers compared to fed livers is indicated above the scatter graph. (C-D) Serum non-esterified fatty acid (NEFA) and triglyceride levels (n = 5-9 per group). The percentage changes in serum levels compared to respective fed control mice are indicated. (E) Representative images of hematoxylin/eosin (H&E) staining of liver sections from *Iqgap1*^*-/-*^ and WT mice (n = 5-6 per group) at 400x magnification. Inset is 800x magnification. (F-G) Hepatic gene expression of *Cpt1a, Acadm, Hadha*, and *Fgf21* normalized to *Gapdh* in WT and *Iqgap1*^*-/-*^mice. (H) Serum FGF21 levels were measured by ELISA (n = 6-7 mice per group).

To characterize the significance of hepatic IQGAP1 induction in fasting, we carefully examined the fasting response in *Iqgap1*^*-/-*^ mice. Upon a 24-hour fast, both WT and *Iqgap1*^*-/-*^mice (31) lost comparable amount of body weight (figure S1B) and liver weight (figure S1C). As expected, serum glucose levels decreased with fasting, but this decrease was exaggerated in *Iqgap1*^*-/-*^ mice since these mice were hyperglycemic in the fed state (figure S1D) and exhibited a trend towards insulin resistance (figure S1E). This is in line with recently published data that showed loss of IQGAP1 protein results in poor glucose tolerance (10). However, the observed reduction in fasting glucose is not due to lack of gluconeogenic response in the *Iqgap1*^*-/-*^ livers (Figure S1F).

Next, we analyzed the effect of IQGAP1 deletion on lipid metabolism. Fasting increased serum non-esterified fatty acids (NEFAs) and modestly reduced serum triglyceride levels in WT animals (figure 1C-D). Even though fasting levels of serum triglyceride and NEFA were significantly lower *in Iqgap1*^*-/-*^ mice, the fold-changes in these metabolites were comparable between the two groups. This is because, compared to WT, the average serum NEFA and triglyceride levels were lower in *Iqgap1*^*-/-*^ mice even in the fed state. To ensure this was not due to poor fat mobilization, we tested the expression of the three main lipases *(Hsl, Lpl*, and *Atgl*) in adipose tissue and found no difference between WT and *Iqgap1*^*-/-*^ (Figure S1G). The *Iqgap1*^*-/-*^ mice lost on average 8% of their visceral adipose by weight upon fasting while the WT mice lost 5%, indicating that lipolysis defects do not contribute to the observed changes in circulating free fatty acid levels (Figure S1H). Furthermore, histological analysis did not reveal any overt difference between WT and *Iqgap1*^*-/-*^ livers, which accumulated triglycerides to a similar extent upon fasting (figure 1C). We then examined the fasting response in the liver by measuring transcript levels of β-oxidation genes Carnitine O-Palmitoyltransferase Ia *(Cpt 1a)*, medium chain acyl-CoA dehydrogenase (*Acadm*), the alpha subunit of hydroxyacyl-CoA dehydrogenase (*Hadha*), *and Fgf21*, a known fasting responsive gene. Despite no difference in induction of *Cpt1a* levels (figure 1F) nor serum ketone body (β-hydroxybutyrate) levels [WT fasting: 1.50 ± 0.36 mM; *Iqgap1*^*-/-*^ fasting: 1.72 ± 0.55 mM], we observed significantly dampened induction of *Acadm, Hadha*, and *Fgf21* in *Iqgap1*^*-/-*^ mice (figure 1G). Consistently, we found that *Iqgap1*^*-/-*^ mice had lower circulating FGF21 levels (figure 1H). These data uncover a role for IQGAP1 in maintaining appropriate hepatic fatty acid metabolism during fasting.

### IQGAP1 is necessary to orchestrate ketogenic adaptation

Since *Iqgap1*^*-/-*^ mice exhibit decreased expression of a few genes involved in β-oxidation and the fasting response, despite remaining capable of mobilizing fat, we examined if long-term ketosis was also altered. To test this, WT and *Iqgap1*^*-/-*^ mice were fed a high-fat, low-carbohydrate ketogenic diet (KD) for 4 weeks. Both cohorts of mice displayed similar overall body weight throughout the experiment. KD feeding resulted in an expected reduction in liver weight, but this response was dramatically blunted in *Iqgap1*^*-/-*^ mice (figure 2A). Histological analysis revealed more microsteatosis in the livers of *Iqgap1*^*-/-*^ mice compared to WT (figure 2B). As a result, their livers appeared paler, and their serum NEFA and triglycerides were significantly lower than WT (figure 2C-D). Further, the gonadal white adipose tissue size, though smaller overall in *Iqgap1*^*-/-*^ mice, increased by a similar proportion in WT and *Iqgap1*^*-/-*^ mice (figure S2A) and showed similar expression of KD-responsive genes (figure S2B) confirming that the metabolic defect is specific to the liver.

**Figure 2.**
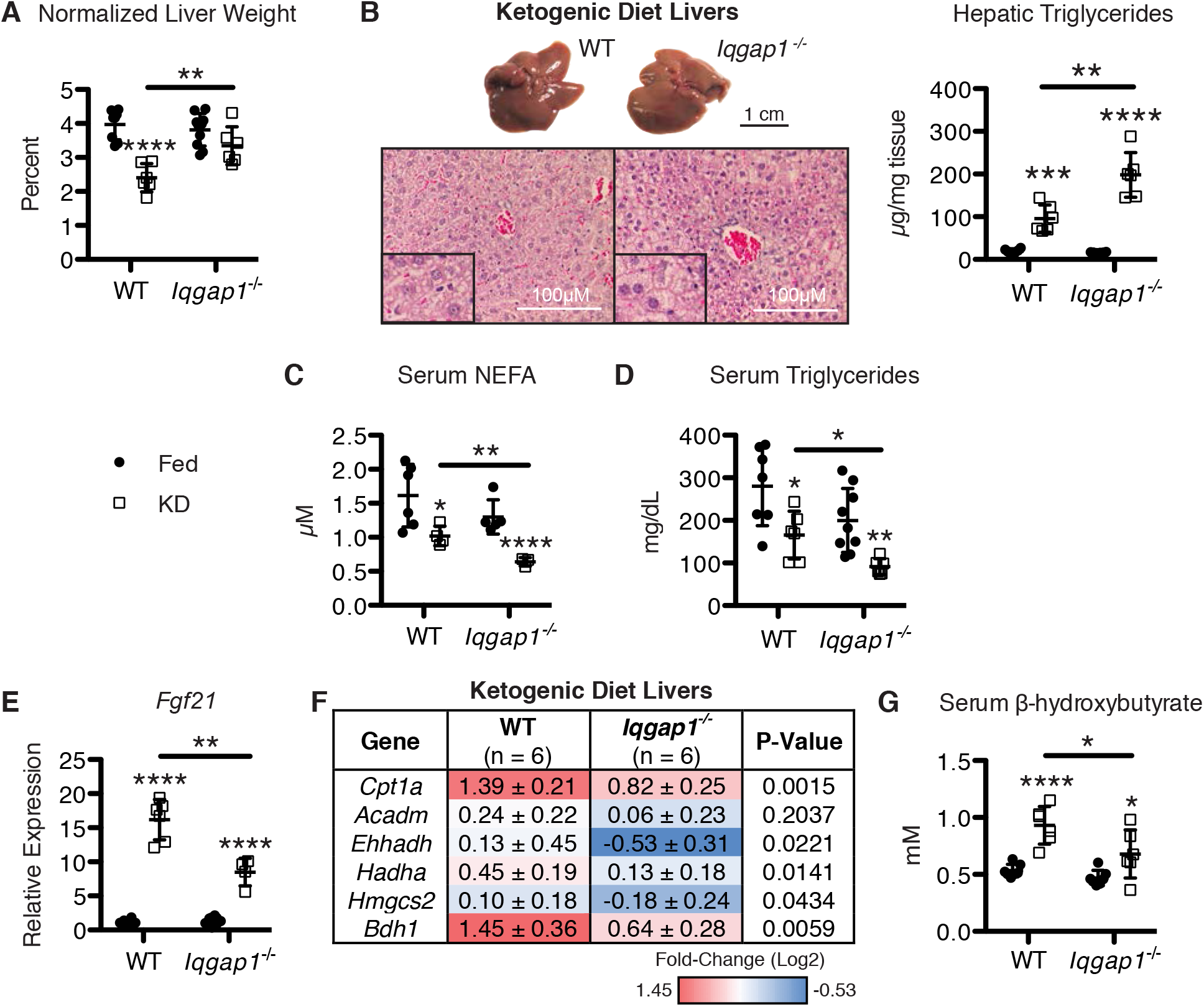
Loss of IQGAP1 results in a blunted ketogenic response. WT and *Iqgap1*^*-/-*^ mice were fed a ketogenic diet (KD) for 4 weeks and fasted overnight (n = 6 per group). As a control, mice were fed normal chow *ad libitum.* (A) Liver weight normalized to total body weight. (B) Gross appearance of livers from KD-fed WT and *Iqgap1*^*-/-*^ mice (*upper*). Representative images of H&E stained liver sections from KD-fed mice (*lower*) at 400x magnification (n = 6 mice per group). Inset is 800x magnification. Hepatic triglyceride was measured in *Iqgap1*^*-/-*^ and WT mice (*right*). (C-D) Serum NEFA and triglyceride levels. (E) Hepatic *Fgf21* expression normalized to *Gapdh* expression in WT and *Iqgap1*^*-/-*^ mice. (F) Heat map summarizing hepatic β-oxidation and ketogenesis gene expression normalized to *Gapdh* expression in KD-fed WT and *Iqgap1*^*-/-*^ mice relative to genotype controls. Values are expressed as mean ± SD. (G) Serum β-hydroxybutyrate levels.

To understand why more fat accumulated in *Iqgap1*^*-/-*^ livers, we examined the hepatic expression of genes controlling fatty acid breakdown and ketogenesis. We found that *Fgf21* (figure 2E), Carnitine palmitatoyltransferase 1A (*Cpt1a*), β-oxidation genes Enoyl-CoA hydratase/3-hydroxyacyl-CoA dehydrogenase (*Ehhadh*), *Acadm*, *Hadha*, and ketogenic genes hydroxymethylglutaryl-CoA synthase 2 (*Hmgcs2*) and 3-hydroxybutyrate dehydrogenase (*Bdh1*), were all significantly down regulated (figure 2F), resulting in lower circulating levels of ketone bodies (figure 2G) in *Iqgap1*^*-/-*^ animals. These results clearly indicate that IQGAP1 is required for long-term adaptation to nutritional ketosis.

### Loss of IQGAP1 deregulates PPARɑ activation subsequent to mTORC1 activation

We next examined whether deletion of IQGAP1 affects the activation of the nuclear receptor PPARɑ, a critical transcriptional regulator of the fasting and ketogenic response (14, 16). We tested PPARɑ downstream signaling in WT and *Iqgap1*^*-/-*^ mice after pharmacological activation with its agonist Wy-14,643 (WY) (32). As expected, WY-treatment induced hepatomegaly in mice. Interestingly, *Iqgap1*^*-/-*^ livers were larger than WT livers (figure S3A). We found that PPARɑ targets including *Cyp4a10*, *Cpt1a, Acadm*, *Ehhadh*, *Hmgcs2*, and *Bdh1* showed significantly reduced induction in *Iqgap1*^*-/-*^ mice while *Hadha* showed a similar but insignificant trend (figure 3A). However, *Fgf21* showed comparable fold induction (figure 3A). Serum ketone bodies also reflected a lower range in *Iqgap1*^*-/-*^ mice (figure S3B). We confirmed the cell autonomous function of IQGAP1 using primary hepatocytes from WT and *Iqgap1*^*-/-*^ mice. When we challenged the primary hepatocytes with WY, only the WT cells were able to increase *Fgf21* transcript levels (figure S3D). These results reveal specific defects in fatty acid β-oxidation and ketone synthesis in *Iqgap1*^*-/-*^ hepatocytes, indicating the requirement of IQGAP1 for these metabolic processes.

**Figure 3.**
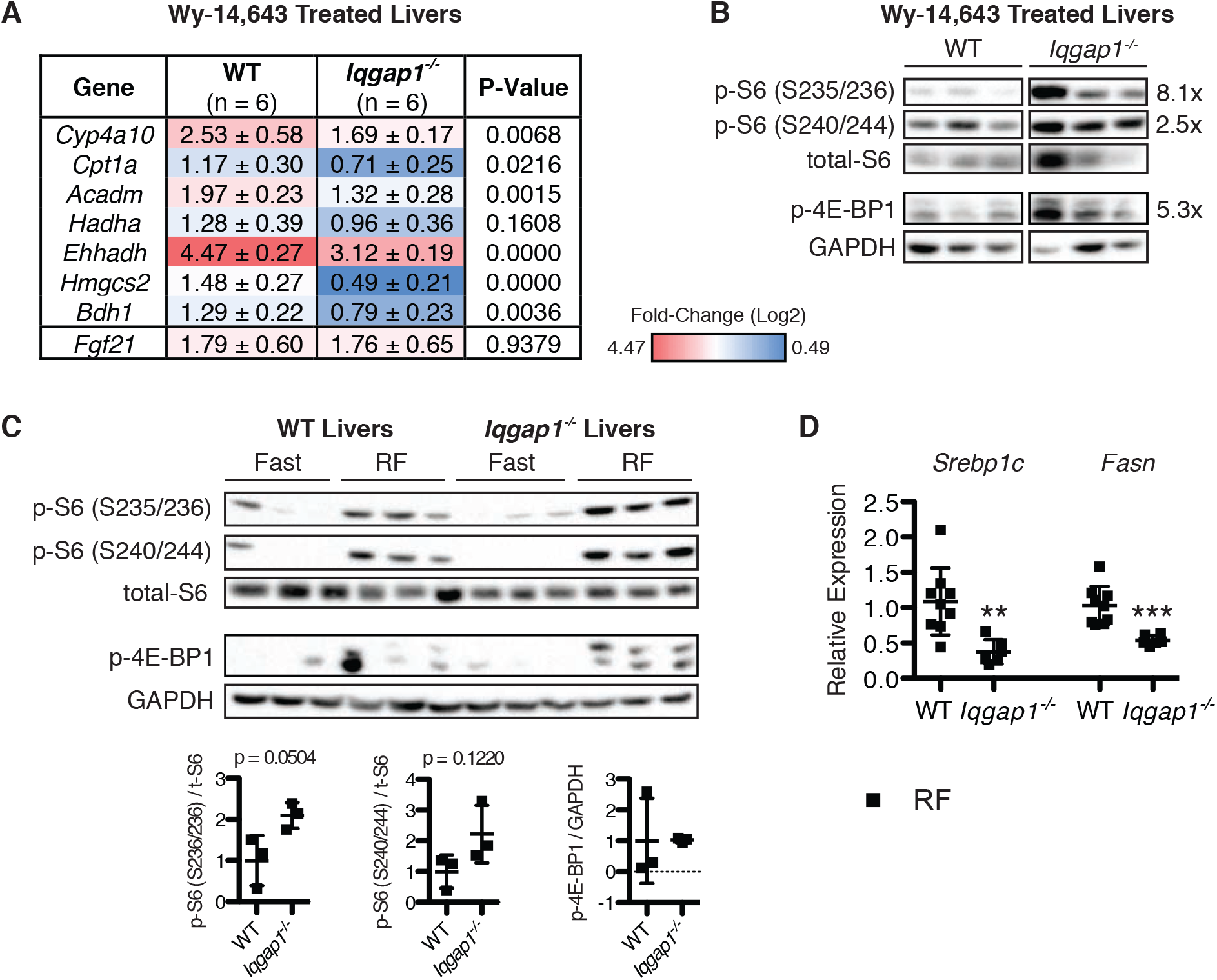
IQGAP1 regulates PPARɑ and mTORC1 activity. (A) Heat map summarizing the fold change in hepatic PPARɑ target gene expression upon WY treatment in WT and *Iqgap1*^*-/-*^ mice relative to CO-treated genotype controls. Expression was normalized to the average of *Gapdh* and *Actin* expression. (B) Immunoblot of liver extracts from WT and *Iqgap1*^*-/-*^ mice treated with WY. Each lane consists of a mixture of liver extracts from two mice (n = 6 mice per group). The amount of phosphorylated-S6 (p-S6) at indicated mTORC1-specific (S240,244) and mTORC1-non-specific (S235,236) serine residues were normalized to total-S6 expression and phosphorylated-4E-BP1 (p-4E-BP1) levels were normalized to GAPDH expression. Relative average expression in *Iqgap1*^*-/-*^ mice compared to WT mice is indicated. (C) Immunoblot of liver extracts from WT and *Iqgap1*^*-/-*^ mice fasted for 24 h or fasted 24 h and refed (RF) 2 h (*upper*). Each lane consists of a mixture of liver extracts from two mice. P-S6 (S235,236) and p-S6 (S240,244) levels were normalized to total-S6 expression and p-4E-BP1 levels were normalized to GAPDH expression (*bottom*). (D) Hepatic gene expression of *Srebp1c* and *Fasn* normalized to *Gapdh* expression in WT and *Iqgap1*^*-/-*^ mice fasted for 24 h and refed 2 h.

It has been previously shown that the major nutrient sensor mTORC1 can suppress transcription of ketogenic PPARɑ targets (21). Therefore, we assessed mTORC1 activity in the liver by measuring phosphorylation of two bona fide markers S6 ribosomal protein and 4E-BP1. S6 phosphorylation at S240,244 and S235,236 sites along with phosphorylation of 4E-BP1 was increased in *Iqgap1*^*-/-*^ livers compared to WT upon WY-treatment (figure 3B). We then examined if the observed elevated mTORC1 activity was unique to WY treatment or was intrinsic in the *Iqgap1*^*-/-*^ livers. To address this, we analyzed fasted and re-fed WT and *Iqgap1*^*-/-*^ mouse livers and found that compared to WT, *Iqgap1*^*-/-*^ mice exhibit a robust 2-fold increase in hepatic mTORC1 activity upon re-feeding (figure 3C). To further confirm increased mTORC1 activation, we analyzed *Srebp1c* and *Fasn* expression levels, which are suppressed in liver-specific TSC1 knockouts that have chronic mTORC1 activation (33). We found robust down regulation of these two genes in *Iqgap1*^*-/-*^ livers (figure 3D). Additionally, we found that phosphorylation of S6 was higher in cultured primary hepatocytes that lacked IQGAP1 (figure S3D), further confirming a cell autonomous role for IQGAP1. To determine if IQGAP1 could regulate mTORC1 through direct interaction, we expressed FLAG-tagged IQGAP1 in HepG2 cells and pulled down IQGAP1-interacting proteins using an anti-FLAG antibody. We found increased pull down of mTOR in cells expressing FLAG-IQGAP1 (figure S3D) implying interaction of IQGAP1 with mTOR. Based on these data we infer that IQGAP1 levels are important in maintaining hepatic mTORC1 activity.

### Hepatic fatty acid oxidation defect in *Iqgap1*^*-/-*^ mice can be alleviated by either re expressing IQGAP1 or inhibiting mTORC1

We re-expressed IQGAP1 specifically in the livers of *Iqgap1*^*-/-*^ mice using an adenoviral system to determine whether this is sufficient for inducing a normal metabolic response. IQGAP1 over-expression resulted in reduction of mTORC1 activity (figure 4A) as well as induction of the fatty acid oxidation genes (figure 4B). Similarly, overexpressing IQGAP1 in the HepG2 cell line reduced S6 phosphorylation (figure S4A). These data reveal that IQGAP1 levels are necessary to maintain mTORC1 signaling. Finally, we tested if inhibiting mTORC1 activation with rapamycin can reverse any of the ketogenic defects observed in *Iqgap1*^*-/-*^ mice. We first ensured that loss of IQGAP1 did not affect mTORC1 inhibition by rapamycin both *in vivo* (figure S4B) and *in vitro* (figure S4C). We found that *Iqgap1*^*-/-*^ mice treated with rapamycin were able to increase a subset of PPARɑ target genes including *Hmgcs2, Fasn*, and *Acadm* to even higher levels than that of the WT (figure 4C). However, *Ppara* and *Cpt1a* levels remained blunted in *Iqgap1*^*-/-*^ mice and was recapitulated in primary *Iqgap1*^*-/-*^ hepatocytes (figure 4D, figure S4C). Overall, these results suggest that hepatic IQGAP1 regulates ketogenesis in an mTORC1-dependent and - independent manner (figure 4E).

**Figure 4.**
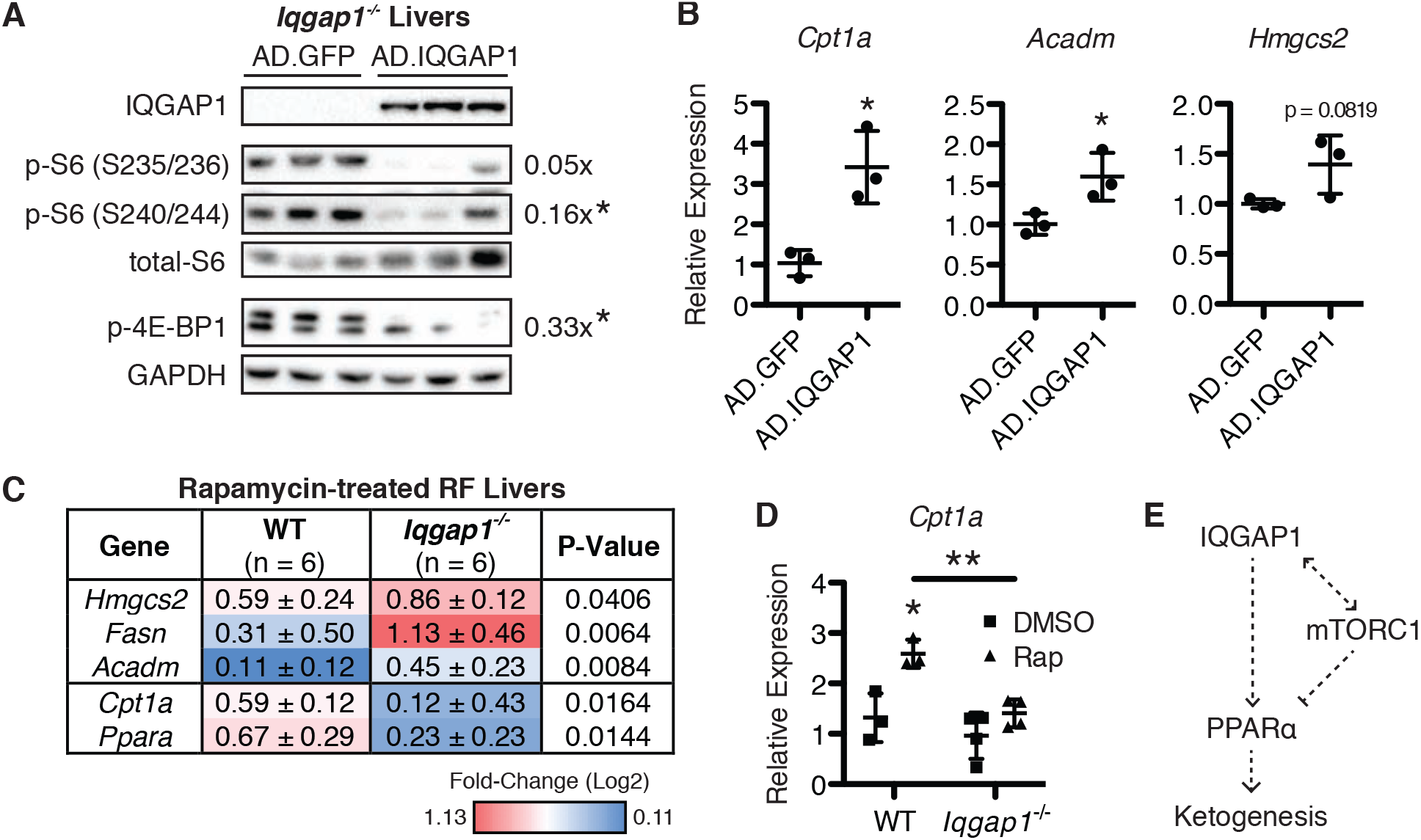
IQGAP1 regulation of a subset of PPARɑ gene targets is mTORC1-dependent. (A) *Iqgap1*^*-/-*^ mice were infected with adenoviruses expressing either GFP or IQGAP1 and sacrificed after 2 weeks. Immunoblot of liver extracts infected with AD.GFP and AD.IQGAP1. Each lane represents an individual mouse (n = 3 mice per group). P-S6 (S235,236) and p-S6 (S240,244) levels were normalized to total-S6 expression and p-4E-BP1 levels were normalized to GAPDH expression. Relative average levels are indicated for AD.IQGAP1 relative to AD.GFP mice. (B) Hepatic gene expression of *Cpt1a, Acadm*, and *Hmgcs2* normalized to *Gapdh* expression in AD.GFP and AD.IQGAP1 infected *Iqgap1*^*-/-*^ mice. (C) WT and *Iqgap1*^*-/-*^ mice were fasted 24 h and refed 2 h (RF) with or without rapamycin (Rap) treatment 1 h before refeeding. Heat map summarizing hepatic fold change in gene expression in Rap+RF mice relative to RF genotype controls. Expression was normalized to *Gapdh* expression. (D) Gene expression of *Cpt1a* normalized to *Gapdh* expression in WT and *Iqgap1*^*-/-*^ hepatocytes treated with 20 nM Rapamycin or vehicle (DMSO) for 6 hours. Each symbol represents hepatocytes from an individual mouse (n = 3-5 per group). (E) Model illustrating how IQGAP1 regulates PPARɑ activation.

## Discussion

IQ motif-containing GTPase Activating Protein 1 (IQGAP1) is a cytoskeleton-associated scaffolding protein that functions in numerous multi-protein complexes and can interact with over 90 proteins. These include the MAP kinase pathway (34), PI3 kinase and AKT (3, 11), calmodulin (35), and FoxO1 (36). To investigate the role for IQGAP1 in coordinating metabolic signaling, we analyzed WT and *Iqgap1*^*-/-*^ mice under different nutritional conditions. We found that IQGAP1 expression was increased upon fasting and *Iqgap1*^*-/-*^ animals showed reduced fatty acid oxidation in the liver. However, increases in gluconeogenic genes were maintained. This could be attributed to intact expression of *Pgc1a* levels, a key regulator of gluconeogenesis (37), in the livers of *Iqgap1*^*-/-*^ mice.

Typically, prolonged fasting leads to ketone body synthesis, which serves an effective alternate fuel source. In particular, we found that IQGAP1 was required to maintain persistent ketogenesis. For instance, when we mimicked this metabolic state using a ketogenic diet (KD) for 4 weeks (38, 39), the *Iqgap1*^*-/-*^ mice showed a poor ability to break down fat and synthesize ketone bodies compared to WT mice. We mapped this to PPARɑ signaling defect in *Iqgap1*^*-/-*^mice. Importantly, *Iqgap1*^*-/-*^ mice mimic *Ppara*^*-/-*^ mice, with both exhibiting elevated hepatic triglycerides, reduced expression of β-oxidation genes, and lower serum FGF21 and ketone body levels during prolonged ketosis (16). Similarly, 24-hour fasting resulted in reduced expression of some genes involved in fatty acid oxidation in *Iqgap1*^*-/-*^ mice relative to WT mice. However, this was not reflected at the serum ketone levels. This is similar to that of the *Fgf21*^*-/-*^ mice, which also show lower ketone body levels when fed KD but do not show changes in ketone levels after 24 hour fast (40, 41). Thus, IQGAP1 may be crucial during long-term fasting specifically.

We confirmed that PPARa downstream signal is dampened when IQGAP1 is deleted by treating mice with the PPARɑ agonist Wy-14,643 (WY). Interestingly, we found that induction of a subset of PPARa target genes responsible for fatty acid oxidation was specifically reduced in *Iqgap1*^*-/-*^ mice. On the contrary, the *Fgf21* response to WY was maintained in the absence of IQGAP1. With *Fgf21* being a starvation hormone, it exhibits dramatic induction after WY treatment in the fasting state (42). Our mice were fed *ad libitum*, which may have potentially contributed to our results. Additionally, it was previously shown that mTORC1 can inhibit PPARa activation (21), and when we analyzed *Iqgap1*^*-/-*^ mice we found increased mTORC1 activity. On the other hand, recent findings (43) show that mTORC1 can positively control *Fgf21* expression. This in turn could explain why *Iqgap1*^*-/-*^ mice do not entirely lack *Fgf21* induction in response to WY. Apart from modulating fatty acid oxidation, WY potently induces liver growth and proliferation in a PPARɑ-dependent manner (44). Contradictively, *Iqgap1*^*-/-*^ mice showed enhanced hepatomegaly compared to WT mice. This intriguing finding in *Iqgap1*^*-/-*^ livers could be secondary to the increased mTORC1 activity, which is a well-known cell growth signal (45). Moreover, we found that IQGAP1 can interact with mTOR in liver cell line indicating a cross talk between the cell growth and scaffolding pathways. In line with our findings, earlier studies have shown that IQGAP1 specifically interacts with mTORC1 (46) and this interaction could promote proliferation (47).

An important finding is that in the liver IQGAP1 levels inversely correlate with mTORC1 activity. Our results demonstrate such that transiently increasing IQGAP1 expression in *Iqgap1*^*-/-*^livers can suppress mTORC1 signaling including S6 and 4E-BP1 phosphorylation. Furthermore, we were able to rescue some of the fatty acid oxidation gene expression in *Iqgap1*^*-/-*^ mice by inhibiting mTORC1 activity with rapamycin. This indicates that mTORC1-independent signals could also contribute towards regulating Cpt1a and PPARa activity in the *Iqgap1*^*-/-*^ mice. Typically, mTORC1 is the major regulator of cellular energetics, but we have not ruled out the role for mTORC2 in this study.

In conclusion, our findings reveal the importance of IQGAP1 in modulating mTORC1 activity and subsequent control of nutritional response in the liver. In addition, PPARa activity, specifically fatty acid oxidation and ketogenesis, is impaired when IQGAP1 is deleted. We therefore postulate that the IQGAP1 scaffold could coordinate multiple facets of long-term ketogenesis.

## Materials and Methods

### Animal experiments

Wild-type (WT) and *Iqgap1*^*-/-*^ mice (Generated in Dr. A. Bernards laboratory and obtained from Dr. Valentina Schmidt, Stony Brook University) maintained on a 129/SVJ background were housed in flow cages at 24 °C on a 12/12 hour light/dark cycle. Genotype was confirmed by PCR analysis of genomic DNA as previously described (31). Male 16-20 week-old mice were used for all experiments. Mice were allowed *ad libitum* access to food and water, except during fasts when they were transferred to a clean cage with access only to water. For refeeding experiments, mice were fasted for 24 h starting at ZT 4 then refed 2 h. A subgroup of the refed mice was injected with 10 mg/kg rapamycin intraperitoneally 1 h prior to refeeding. Their standard chow diet was Teklad F6 Rodent Diet (8664, Envigo), consisting of 31%, 19%, and 50% kcal from protein, fat, and carbohydrate, respectively. In contrast, the Ketogenic Diet (TD.96355, Envigo) consisted of 9.1%, 90.5%, and 0.4% kcal from protein, fat, and carbohydrates, respectively. For fibrate treatment, Wy-14,643 (WY, Cayman Chemical) was dissolved in 100% ethanol to a final concentration of 16.5 mg/mL. The WY solution or vehicle was further diluted with corn oil (CO) to a final concentration of 5 mg/mL and stirred under vacuum overnight to remove excess ethanol. Mice were administered 0.01 mL of CO with or without WY per gram of body weight (50 mg/kg) by oral gavage for four consecutive days and sacrificed on the fifth day. For hepatic expression of GFP and IQGAP1, female *Iqgap1*^*-/-*^ mice were administered high-titer adenovirus expressing either cDNA via tail vein injection. Three days after injection, mice were started on a 0.5 g/kg doxycycline (dox) diet and sacrificed after 2 weeks. All mice were sacrificed at ZT 4-6. Zeitgeber (ZT) time refers to a 24-hour light/dark cycle, with ZT0 corresponding to the appearance of light. Blood was collected from each mouse by retro-orbital bleeding just prior to sacrifice. Serum was separated by centrifugation and stored at - 80 °C in opaque tubes. Liver and gonadal white adipose tissues were collected and flash frozen for analysis. A portion of each tissue was also fixed in 10% formalin for histological analysis. All animal studies were approved by the University of Illinois at Urbana-Champaign Institutional Animal Care and Use Committee and were carried out as outlined in the Guide for the Care and Use of Laboratory Animals prepared by the National Academy of Sciences and published by the National Institutes of Health.

### Primary hepatocyte culture

Mouse hepatocytes were isolated using a two-step collagenase perfusion technique (48). Briefly, male adult 129/SVJ WT and *Iqgap1*^*-/-*^ mice were perfused with 50 mL of Solution 1 (1 μM EDTA in 1x Hanks Balanced Salt Solution without Ca^2+^ or Mg^2+^). Livers were then perfused with 50 mL of Solution 2 (3000 U of collagenase type I from Worthington, 0.54 μM CaCl2, 40 μg/mL Trypsin Inhibitor, and 15 mM HEPES pH 7.4 in 1x Hanks Balanced Salt Solution with Ca^2+^ and Mg^2+^). The perfused liver was then transferred to a Petri dish containing wash buffer (Williams E media with 1x Penicillin/Streptomycin and 1x L-Glutamine) and gently massaged to obtain loose cells. The cell suspension was filtered through a 70 μm filter and centrifuged at 600 *x g* for 4 min. The pellet was suspended in 25 mL of wash buffer and layered onto a Percoll gradient and immediately centrifuged at 600 *x g* for 10 min. The pellet, enriched for live hepatocytes, was washed 3 times then cells were suspended in growing medium (wash media supplemented with 1x Insulin-Transferrin-Selenium (ITS) solution from Gibco) and plated at 5 x 10^5^ live cells/mL in 6-well collagen-coated plates. Media was changed 4-6 hours after plating to remove dead cells. Cells were either cultured for another 24 hours in growing medium or cultured in media without ITS overnight and treated with 20 nM rapamycin in the presence of 100 nM insulin for 6 hours.

### Adenovirus production and cell culture

HepG2 cell line was obtained from ATCC (Catalogue no. HB-8065) and cultured according to ATCC specifications. This cell line tested negative for mycoplasma (Biotool, catalogue no. B39032). For *in vivo* infection, adenoviruses expressing full length *Iqgap1* and *Gfp* were generated as previously described (49). For IQGAP1 overexpression in cell culture, HepG2 cells were cultured in tetracycline (tet)-free DMEM and transfected with tet-inducible FLAG-tagged *Iqgap1* and rtTA plasmids using Mirus TransIT-X2 kit. The next day, the media was replaced with fresh DMEM containing 2 ng/mL dox, and protein was collected 24 hours later.

### Serum chemistry

Serum was thawed on ice prior to each metabolite assay. Assays were run in duplicate for triglycerides (Infinity Triglyceride Stable Reagent, Fisher Scientific), β-hydroxybutyrate (β-hydroxybutyrate (Ketone Body) Colorimetric Assay Kit, Cayman Chemical), and free fatty acids (Free Fatty Acid Fluorometric Assay Kit, Cayman Chemical). All assays were performed according to the kit instructions, unless stated otherwise. For the Free Fatty Acid Kit, absorbance was read at 570 nm and was adjusted for background reading at 600 nm. Additionally, Mouse/Rat FGF-21 Quantikine ELISA kit (R&D Systems) was used to measure serum FGF21 levels. Serum glucose was measured using one-touch glucose strips on fresh tail blood.

### Glucose tolerance test

Glucose tolerance test (GTT) was performed as previously described (50). Briefly, mice were fasted for 13 hours overnight and administered 2g/kg D-glucose intraperitoneally. Blood glucose concentrations were measured at 0, 15, 30, 60, and 120 min after injection using tail blood.

### Hepatic triglyceride assay

Frozen liver tissue (<50 mg) was weighed and homogenized in 1 mL of isopropanol. Samples were centrifuged at 4°C and 10,000 rpm for 15 minutes, and supernatant-containing triglycerides was separated. Triglyceride content was measured using Infinity Triglyceride Liquid Stable Reagent (Fisher Scientific).

### Histology

Formalin fixed liver samples were embedded in paraffin wax. Five micron sections were cut and used for hematoxylin and eosin (H&E) staining according to standard methods (51).

### Quantitative RT-PCR

Total RNA from frozen whole liver tissue was extracted using TRIzol solution (Invitrogen) according to manufacturer’s protocol. RNA quality was determined by A260/280 and bleach RNA gel as previously described (52). RNA (5 μg) was treated with DNase (Promega) and reverse transcribed using random hexamer primer (New England Biosciences) and Maxima Reverse Transcriptase kit (Thermo Scientific). The cDNA was diluted to 12.5 ng/μL with molecular grade water (Corning) and used for qRT-PCR assays. qRT-PCR was performed on an Eco Real-Time PCR system (Illumina) in triplicate using 50 ng of cDNA per reaction and PerfeCTa SYBR Green FastMix (Quanta). All assays were run with an initial activation step for 10 min. at 95 °C, followed by 40 cycles of 95 °C for 10 sec and 60 °C for 30 sec. Primer sequences are described in Table S1. *Gapdh*. *β-Actin*, and *36b4* were used as housekeeping genes.

### Western blot analysis

Protein was extracted from approximately 50 mg of frozen liver tissue. RIPA buffer (25mM Tris pH 7, 150mM NaCl, 0.5% sodium deoxycholate, protease inhibitor (Pierce Protease Inhibitor Mini Tablets, EDTA Free; ThermoFisher Scientific), phosphatase inhibitor (Phosphatase Inhibitor Cocktail 3; Sigma Aldrich), and 1% Triton X-100) was added to each liver sample, which was homogenized by adding 8-10 1mm beads and bullet blending for 3 x 1 minute with 1 minute on ice in between. Samples were sonicated until non-viscous. Protein concentration was measured by BCA assay (Pierce BCA Protein Assay Kit, Thermo Fisher Scientific). For western blot, 10-50 μg of total protein was loaded onto 8-12% SDS-PAGE gels. After transfer, the membrane was incubated with antibodies described in Table S2.

### Co-Immunoprecipitation

HepG2 cells were transfected with dox-inducible FLAG-IQGAP1 and were treated with dox or vehicle for 24 hours. After induction, approximately 10^7^ cells were lysed in lysis buffer (50 mM Tris pH7.8, 150 mM NaCl, 30mM EDTA, 0.5% Triton X100). The lysate was then split and equal amounts were immunoprecipitated with a FLAG antibody overnight, washed, and resolved on an 8% SDS-PAGE gel. After transfer, blots were probed with antibodies for mTOR.

### Statistical Analysis

Data are expressed as mean ± SD. Student’s unpaired two-tailed t-test was used to compare two groups. A * indicates a statistically significant difference between the treatment group and its respective genotype control unless the groups are otherwise indicated. Significance is defined as p < 0.05, with * p < 0.05, ** p < 0.01, *** p < 0.001, and **** p < 0.0001. Outliers were determined by Grubbs’ test and removed from analysis.

## Acknowledgements

Authors would like to thank Drs. Andre Bernards at MGH, Boston and Valentina Schmidt at Stony Brook University, New York for sharing the *Iqgap1*^*-/-*^ mice. We also like to thank Dr. Jie Chen and Dr. Shomit Sengupta for their critical comments and suggestions during this study. We thank Ms. Karen Wendt who initiated the fasting experiments in *Iqgap1*^*-/-*^ mice. This study was supported by start up funds from University of Illinois at Urbana-Champaign (S.A) and HD080011 NICHD (S.A).

## Author Contributions

H.L.E, and S.A. contributed to the conception, design, data acquisition, analysis and drafting of the manuscript.

